# A CLEM approach to access to the ultrastructure at the graft interface in *Arabidopsis thaliana*

**DOI:** 10.1101/2021.07.13.452258

**Authors:** Clément Chambaud, Sarah Jane Cookson, Nathalie Ollat, Emmanuelle Bayer, Lysiane Brocard

## Abstract

Despite recent progress in our understanding of the graft union formation, we still know little about the cellular events underlying the grafting process. This is partially due to the difficulty of reliably targeting the graft interface in electron microscopy to study its ultrastructure and three-dimensional architecture. To overcome this technological bottleneck, we developed a correlative light electron microscopy approach (CLEM) to study the graft interface with high ultrastructural resolution. Grafting hypocotyls of *Arabidopsis thaliana* lines expressing YFP or mRFP in the endoplasmic reticulum allowed the efficient targeting of the grafting interface for under light and electron microscopy. To explore the potential of our method to study sub-cellular events at the graft interface, we focused on the formation of secondary plasmodesmata (PD) between the grafted partners. We showed that 4 classes of PD were formed at the interface and that PD introgression into the call wall was initiated equally by both partners. Moreover, the success of PD formation appeared not systematic with a third of PD not spanning the cell wall entirely. Characterizing the ultrastructural characteristics of these failed PD gives us insights into the process of secondary PD biogenesis. We showed that the thinning of the cell wall and the endoplasmic reticulum-plasma membrane tethering seem to be required for the establishment of symplastic connections between the scion and the rootstock. The resolution reached in this work shows that our CLEM method offer a new scale to the study for biological processes requiring the combination of light and electron microscopy.

## Introduction

Grafting plants permits us to select and combine different rootstock and scion phenotypes, such as root pathogen resistance and fruit quality traits, independently of genetic introgression (Mudge, 2009). Nevertheless, graft compatibility is an essential prerequisite (Flaishman et al., 2008; Chen et al., 2017). As such, the study of graft union formation and the characterization of the mechanisms involved are important for the breeding of many horticultural crops (Kyriacou et al., 2017). In addition, grafting is also used in scientific studies to study long-distance movement of molecules and to graft-inoculate pathogens (Ragni et al., 2011; Liang et al., 2012; Bidadi et al., 2014; Zhang et al., 2014; Simone et al., 2016; Ota et al., 2020).

Grafting involves cutting the scion and rootstock, and maintaining the cut surfaces in close proximity. During grafting, as opposed to the wounding, all physical cell communications are interrupted by cutting at the grafting point. To survive, both grafted partners, have to interact together quickly and effectively to establish new connections. During graft union formation the cells at the interface collapse, a necrotic layer forms, and the scion and rootstock adhere to each other (Sala et al., 2019). Cells then proliferate at the cut surfaces to form a callus, some of these callus cells are thought to differentiate into phloem and xylem that connect the vasculature of the scion and rootstock (Melnyk et al., 2015). Recent progress has been made in understanding of graft union formation by using the *Arabidopsis thaliana* hypocotyl grafting model (Matsuoka et al., 2016; Sala et al., 2019; Tsutsui et al., 2020). In this model, the developmental framework and transcriptomic changes have been excellently documented (Melnyk et al., 2015; Melnyk et al., 2018). The establishment of a vascular continuum begins by phloem reconnection at 3 days after grafting (DAG) and then, xylem reconnection occurs from 5 to 7 DAG (Melnyk et al., 2015). New vascular tissue formation is initiated above, and then below the graft interface indicating that the scion could be the leader of this cellular differentiation processes. At the graft interface, new differentiated vascular cells expand and divide to bypass the wounded regions and permit the vascular reconnection between both partners (Melnyk et al., 2015).

While long-distance signalling is ensured by the vascular tissues, cell-to-cell communication is mediated via nanometric channels called plasmodesmata (PD) (Li et al., 2021b). These channels span the cell wall and participate in developmental processes through the trafficking of cytosolic and membrane molecules between neighbouring cells (Cantrill et al., 1999; Lucas and Lee, 2004; Benitez-alfonso et al., 2009; Stonebloom et al., 2009; Xu et al., 2011; Furuta et al., 2012). Plasmodesmata are composed of a central cylindrical element, called the desmotubule (Tilney et al., 1991; Ding et al., 1992). The desmotubule, localized exactly at the cell-to-cell interface, is a specialized constricted domain of endoplasmic reticulum (ER) shared between both cells. Inside the PD, the desmotubule interacts physically with a particular plasma membrane domain, which is in continuity with the plasma membranes of neighbouring cells (Grison et al., 2015; Nicolas et al., 2017; Brault et al., 2019). The space between the desmotubule and the plasma membrane, called the cytoplasmic sleeve, is not visible in very young PD (PD of type I, i.e. PD with no discernible cytoplasmic sleeve), but is larger in more mature PD (PD of type II, i.e. PD with visible space between the desmotubule and the plasma membrane) (Nicolas et al., 2017). Another nomenclature used to describe PD, primary and secondary PD, distinguishes PD based on their biogenesis (Ehlers and Kollmann, 2001). Primary PD are formed during cell division by trapping ER strands in a neo cell wall during its formation (Hepler, 1982; Nicolas et al., 2017). Whereas secondary PD are formed in pre-existed cell walls by an unknown mechanism.

The studies of Kollmann and Glockmann (Kollmann et al., 1985; Kollmann and Glockmann, 1985; Kollmann and Glockmann, 1991) on the heterograft *Vicia faba/Helianthus annuus,* where callus cells possess species-specific ultrastructural differences, demonstrated the existence of PD at the graft interface. Because the scion and rootstock cells belong to different cell lineages, PD at the graft interface are necessarily secondary. Unfortunately, this approach is limited to studying very different species grafted together, so possibly characterises grafts with a certain degree of incompatibility (at best only 50 % of *V. faba/H. annuus* grafts survived 4-6 weeks) and does not necessarily give insights into the development of compatible grafts. In addition, relying on species-specific ultrastructural differences does not allow the unambiguous identification of every cell studied because of the intrinsic ultrastructural cell heterogeneity occurring between different cells.

Horizontal-DNA transfers have been found at the graft interface of tobacco grafts (Stegemann and Bock, 2009; Stegemann et al., 2012; Hertle et al., 2021), suggesting that large openings form at the graft interface. A recent study showed that callus cells of the graft interface could have cell wall holes of approximately 1.5 μm in diameter, which could allow DNA and organelle transfers (Hertle et al., 2021). Despite an attempt made to track plastids with live-imaging to show plastid exchanges through these holes, the absence of a complete three-dimensional (3D) view of the observed cells and of a single plastid tracking method means that the authors could not unequivocally show that movements of a plastid observed around a cell wall correspond to the same plastid going through the cell wall (Hertle et al., 2021). Therefore, direct evidence of the movement of organelles between the scion and rootstock of grafted plants has yet to be found.

In order to gain insights into the ultrastructural events underpinning the graft union formation, we developed a Correlative Light and Electron Microscopy approach (CLEM) on grafted hypocotyls of *A. thaliana* to target the graft interface. Correlative light and electron microscopy approaches have been developed for several model organisms and have been started to use for plant tissues (Bell et al., 2013; Marion et al., 2017; Wang and Kang, 2020). The different used protocols published to date give access to the ultrastructure context of fluorescence labelling, but the ultrastructure preservation and resolution are limited to relatively low magnifications and provide only ultrastructural overviews. Here, we describe in detail a CLEM protocol to localize and study for the first time with a very high reliability the graft interface of the model plant *A. thaliana.* This method is based on the in-resin fluorescence CLEM methods published by Kukulski et al., 2012 that we adapted for plant samples. Combined with electron tomography (ET), it permits us to study the fine 3D ultrastructural details of our samples, up to the resolution of the bilayer of the plasma membrane. Fluorescent proteins were used to label the scion and the rootstock, and permitted us to localize the graft interface under light microscopy before revealing its ultrastructure with TEM and ET. The ability to identify the origin and the type of cells at the graft interface offer new perspectives to understand of cytological events involved in establishing communication between the scion and rootstock of grafted plants. We chose as a proof of concept to use this approach to answer the following questions: Are PD formed at the graft interface of homo-grafted hypocotyls? Do the PD formed at the graft interface differ in ultrastructure? What are the characteristics of successful PD biogenesis at the graft interface and how can they inform us on the mechanisms involved? Do both the scion and rootstock initiate PD biogenesis, if so, are both partners equally involved?

## Materials & Methods

### Plant material, growth conditions and plant grafting

*Arabidopsis thaliana* were grown vertically in a growth chamber on solid medium composed of half Murashige and Skoog medium including vitamins (1/2 MS) and plant-Agar (8 g L^-1^), pH 5.8. Growth conditions were set at 22 °C in a growth chamber with a 10 h photoperiod with photosynthetic photon flux density of 120 μmol m^-2^ s^-1^. All lines (35S::HDEL_mRFP, 35S::HDEL_YFP and 35S::GFP) were previously published (Nelson et al., 2007; Lee et al., 2013; Clark et al., 2016). 35S::TagRFP were produced in a Col-0 background.

For grafting, seven-day-old plants were selected and grafted using the transverse cut and butt alignment method described by Melnyk (Melnyk, 2017). This method was slightly modified: ultra pure water plus plant-Agar (1.6 g L^-1^) was used instead of humid paper to maintain the moisture level required for graft union formation.

### High Pressure Freezing (HPF) and Cryosubstitution

Copper carriers (100 μm deep and 1.5 mm wide, Leica Microsystems, catalog number 16707898) were filled with ultra pure water plus 20 % bovine serum albumin (BSA) (which functions as a cryoprotectant). Rapidly, grafts at 3 or 6 DAG were cut above and below the graft point and installed in the well of the carrier. Samples were frozen in the carriers using an EM-PACT high-pressure freezer (Leica) and then transferred at −90 °C into an AFS 2 freeze-substitution device (Leica). Grafts were incubated in a cryosubstitution mix containing only uranyl acetate 0.1 % in pure acetone for 30 h (freeze substitution and embedding protocol adapted from Nicolas et al., 2018). Afterwards temperature was raised progressively to −50 °C, at 3 °C h^-1^. The cryosubstitution mix was removed and thoroughly washed 3 times with pure acetone and then 3 times in pure ethanol. Low temperature and the limited amount of chemical compounds help in preserving the fluorescence of FPs while preserving a sufficient staining for TEM and ET. Samples were then carefully removed from the carriers before progressive embedding in HM20 Lowicryl resin (Electron Microscopy Science): HM20 25 % 2 h, 50 % 2 h, 75 % overnight (diluted in pure ethanol), HM 20 100 % 2 h twice before a last 100 % for 8 h 25 %, 50 % (2 h each), 75 % (overnight), 100 % (twice for 2 h) and a last 100 % for 8 h. The polymerization was done under UV light for 24 h at −50 °C followed by 12 h at +20 °C. Samples in resin blocks were stored at −20 °C protected from light.

### Microscope acquisitions and correlations

Prior to observation, sections were made with an EM UC7 ultramicrotome (Leica). For an optimal balance between resolution under the electron microscope and signal quantity under the confocal microscope, sections from 150 nm to 200 nm were collected on parlodion coated 200-square mesh copper grids with thin bars (Electron Microscopy Science). Fluorescence Z-stack acquisitions were made on a Zeiss LSM 880 with excitation 488 nm for YFP and 561 nm for mRFP. The acquisitions were achieved with a 63× apochromatic N.A 1.4 oil objective. The tile acquisition tool of the Zen black software was used to get the large field of view with the best resolution. Moreover, the Z-stack tool was use to acquire the maximum of photons emitted by the samples. Afterwards maximal or summed Z-projections were used to improve the visualization. Transmission electron microscopy observations were carried out on a FEI TECNAI Spirit 120 kV electron microscope equipped with an Eagle 4Kx4K CCD camera. Correlation was based on natural landmarks such as cell shapes, plastids and nucleus. Correlated images were obtained thanks to the plugin ec-Clem (Paul-Gilloteaux et al., 2017) of Icy software (De Chaumont et al., 2012).

### Tomogram acquisition and reconstruction

We used the process described by (Nicolas *et al.,* 2018). Before tomogram acquisition, sections need to be coated with gold fiducials for subsequent alignment, a 1:1 mixture of 0.5 % BSA and 5 nm colloidal gold solution from BBI solutions (referenced as EM-GC5 on http://bbisolutions.com) was used. Tilt series were acquired with a single tilt specimen holder (Fichione instruments, model-2020), using the batch mode of FEI 3D Explore tomography software. Each tilt series was acquired between −65° to 65° when it was possible, with an acquisition at every degree. For dual axis, the grid was turned to 90° manually. The raw tilt series were aligned and then reconstructed using the fiducial alignment mode with the eTomo software (http://bio3d.colorado.edu/imod/). 10 to 25 fiducials were used to accurately align all images. Reconstruction was performed using the back-projection with SIRT-like filter (10 to 50 iterations).

### Image processing and analysis

Image analysis was carried out using the ImageJ software (https://imagej.nih.gov/ij/download.html) with the Bio-Format plugin (https://www.openmicroscopy.org/bio-formats/) to open and process the Zeiss .czi files and to open the .mrc tilt series files and tomograms. Measurement of cell wall thickness was done using ImageJ. Mosaic images from the confocal microscope were assembled with the BigStitcher plugin of Fiji. Mosaic images of TEM were assembled using Photomerge function of Adobe Photoshop. No image analysis and measurements were carried out on mosaic and correlated images, they were only used as a visualization tools.

### Data analysis

The thickness of cell walls around PDs was done by measuring the cell wall thickness at 5 locations distributed in the vicinity of PD followed by the calculation of the mean thickness of the cell wall around each PD. Data analysis and box plot presentation were done using SigmaPlot, Systat Software Inc.

## Results

### 1. Targeting the graft interface using Correlative Light Electron Microscopy

In order to precisely and accurately identify the graft interface of hypocotyls of *A. thaliana* at the cellular level, we used green and red intracellular FPs to tag the scion and rootstock respectively. To do this we made use of transgenic lines of *A. thaliana* expressing either YFP (yellow FP) or mRFP (monomeric red FP) with an ER retention signal (HDEL_YFP or HDEL_mRFP) (Fig. 1A and B). The HDEL retention signal prevents the FPs from being secreted and retains them in the ER lumen (Pelham et al., 1988). We visualized graft interfaces of the scion/rootstock combination HDEL_YFP/HDEL_mRFP by confocal microscopy at 5 DAG; we observed a clear demarcation between YFP- or mRFP-containing cells at the graft interface (Fig. 1B). As a positive control, we used *A. thaliana* lines expressing a free cytosolic FP (GFP or mRFP), which are known to diffuse from cell-to-cell through PDs. When the lines expressing a free cytosolic FP were used as one graft partner, the cells in the vicinity of the graft interface were systematically labelled with both the cytosolic and ER-bound FP (Fig. 1 C and D). We concluded that, as expected, HDEL tagged FPs cannot freely pass from one grafted partner to the other and that using the HDEL_YFP and HDEL_mRFP expressing lines permits the precise and accurate identification of the graft interface.

**Figure 1.**
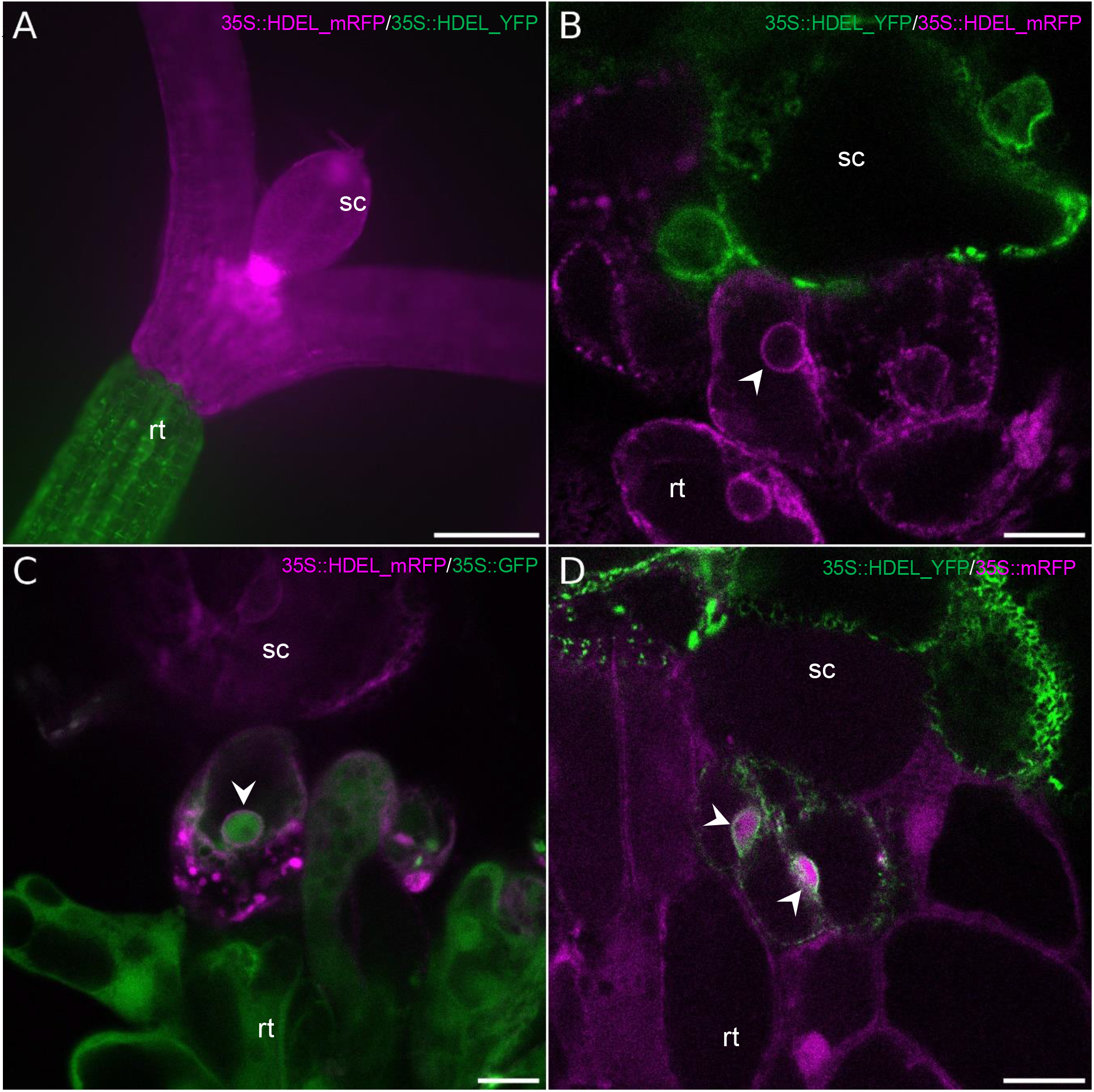
Targeting the graft interface of *Arabidopsis thaliana* hypocotyl grafts with fluorescent proteins. The scion (sc) and the rootstock (rt) are at the top and bottom positions respectively. **(A)** Large view of a graft between a scion expressing 35S::HDEL_mRFP (red) onto a rootstock expressing 35S::HDEL_YFP (green). **(B)** Observation by confocal microscopy of a graft interface of 35S::HDEL_YFP (scion) / 35S::HDEL_RFP (rootstock). No YFP nor RFP signal is found in the opposite partner. **(C)** Observation by confocal microscopy of a graft interface of 35S::HDEL_RFP (scion) / 35S::GFP (rootstock). GFP and HDEL-mRFP can be observed in same cells. **(D)** Observation by confocal microscopy of a graft interface of 35S::HDEL_YFP (scion) / 35S::mRFP (rootstock). mRFP and HDEL-YFP can be observed in same cells. White arrowheads pinpoint nuclei. Scale bars: (A) 200 μm, (B, C and D) 10 μm

To investigate the graft interface at the ultrastructural level, we then used a CLEM approach on the grafted HDEL_YFP/HDEL_mRFP hypocotyls (Fig. 2). The graft interfaces could be harvested from 3 DAG without the scion and rootstock detaching from each other. After harvesting, graft interfaces were then quickly fixed by HPF in order to ensure the biological preservation of samples closest to the native state. During freeze substitution steps, the fluorescence preservation of YFP and mRFP needs to be maintained without compromising the ultrastructure preservation. To this end, fixative contrasting compounds were limited to uranyl acetate and we used limited incubation times for acetone, resin and UV exposition. Following freeze substitution in HM20 resin, ultrathin sections of 70 nm were made and harvested on TEM grids and then visualized under confocal microscopy after mounting in water between a glass slide and a cover slip. Despite the preventive measures during fixation, the YFP and mRFP fluorescence signals were too weak to discriminate with enough confidence a FP-specific signal. Moreover, the formation of air bubbles between the section and the cover slip had the effect of increasing refraction phenomena and led to the loss of an important part of emitted photons (data not shown). To overcome this technical hurdle, the thickness of ultrathin sections was increased from 70 nm to 150 nm to maximize the total number of FPs within a single section. Precautions were made during sample mounting to permit the complete water immersion of the section by maintaining the grid on a drop of water with a thin pin during the positioning of the cover slip. In order to improve the visualization of FPs and because the TEM grids were not perfectly flat, Z-stack acquisitions followed by maximal Z-projections were systematically performed. Overall, YFP detection required a higher excitation power and a numerical compression of the intensity levels compared to mRFP. Thus, mRFP fluorescence is less affected during TEM sample preparations than YFP (Fig. S1). We also noted that the TEM-sample-embedding process induced a shift for chlorophyll emission spectra that overlaps with mRFP spectra (Fig. S2). Despite this, cells expressing mRFP and YFP ER-markers were reliably differentiated and graft interfaces were targeted without ambiguity (Fig. 2). Following confocal acquisition, the TEM observation of the very same section gave access to the ultrastructural data. Correlation between both microscopies, based on the overlay of natural markers (the cell shapes, plastids and nuclear envelops), revealed with no ambiguity the identity and the ultrastructure of each cells with one exception for complete differentiated TE and SE. In fact, complete differentiation process for SE and TE result in loss or modification of ER, which was not still labelled with HDEL-markers. Despite this loss of labelling, correlation permitted to identify the vascular tissues especially at the graft interface and access to the SE and TE continuities between both grafted partners (Fig. 2). Magnified observations permitted us to evaluate the level of ultrastructural preservation. The cell compartmentation was conserved with notably the maintenance of the integrity of chloroplasts, nuclei, Golgi and vesicles (Fig. 2G & H). An important technical advance, compared to previous CLEM protocol in plants (Bell et al., 2013; Marion et al., 2017; Wang and Kang, 2020), is the capacity to reveal 3D shapes with high ultrastructural details thanks to ET. The different parts of PD, such as the bilayer of plasma membrane and the desmotubule were spatially resolved indicating the high quality of the ultrastructure preservation despite the absence of strong chemical fixative agents (Fig. 2I). Moreover, our method permits to preserve the antigenicity of the sample and is compatible with immuno-gold labelling (data not shown).

**Figure 2.**
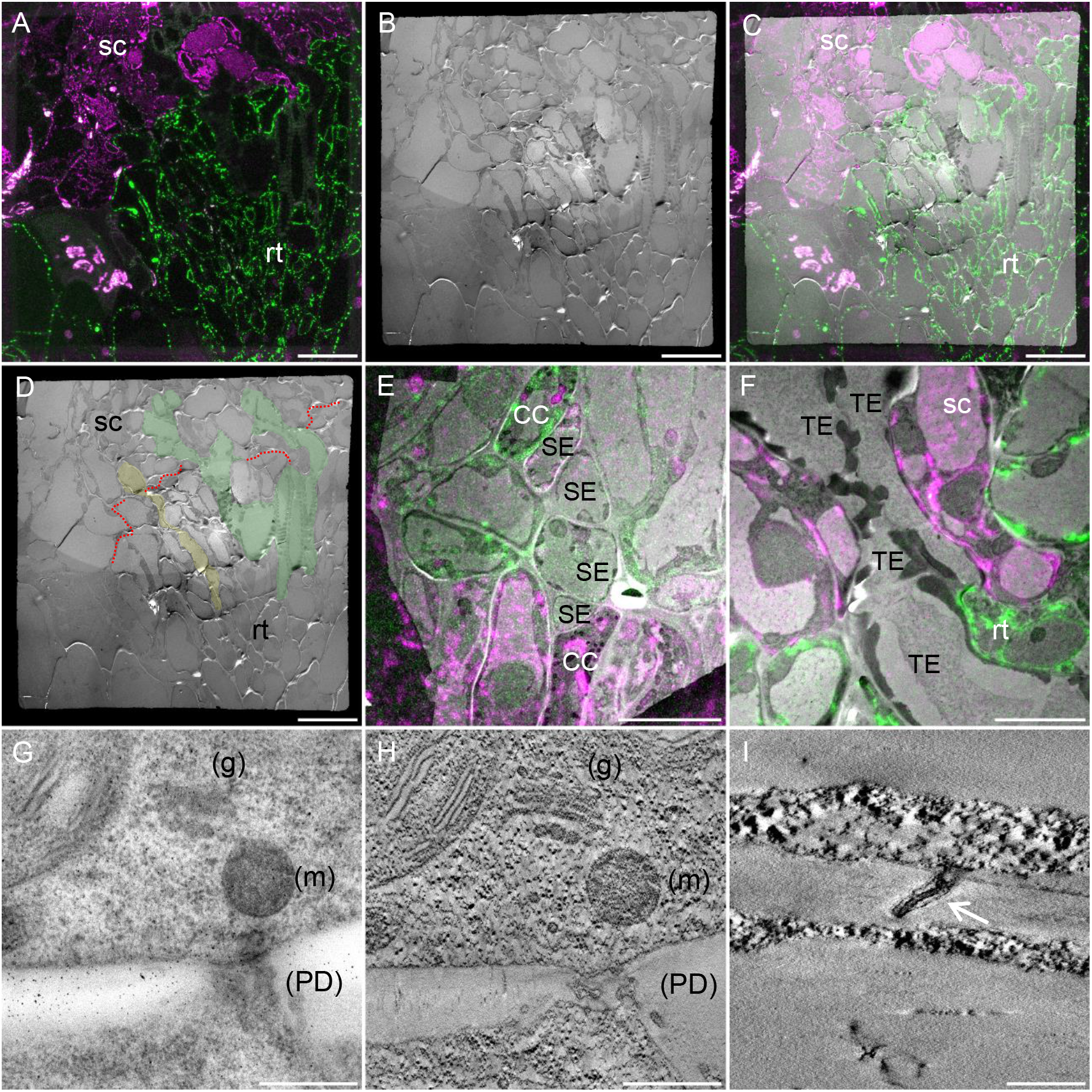
Correlative Light Electron Microscopy as a tool to investigate the ultrastructure of the graft interface. The scion and the rootstock are respectively at the top and bottom positions (A, B, C, E and F). **(A)** Confocal acquisition of the graft interface of a 35S::HDEL_mRFP scion grafted onto a 35S::HDEL_YFP rootstock **(B)** Transmission electron micrograph tiles of the regions observed in (A). **(C)** Correlated light electron microscopy view of the graft interface which allows the precise identification of the scion and rootstock cells. **(D)** Thanks to correlative images, the “map” of the graft interface (red dotted line) is made and allows us to identify the scion (sc), the rootstock (rt), phloem vessels (yellow) and xylem vessels (green). **(E)** Correlated view of sieve elements (SE) and compagnion cells (CC) aligned at the interface. **(F)** Correlated view of Xylem tracheary element (TE) properly aligned at the graft interface. **(G)** Electron micrograph of a cell wall shows the golgi apparatus (g), mitochondria (m), and plasmodesmata (PD) **(H)** its corresponding tomogram the increase in resolution shows the preservation of the membranes of the organelles. **(I)** Tomogram showing the plasmodesmata plasma membrane (white arrow) and the desmotubule membranes. White arrowheads denote “natural landmarks” i.e. plastids, mitochondria, nuclei, that were used to correlate confocal and transmission electron microscopy images. Scale bars: (A, B, C and D) 20 μm, (E) 10 μm, (F) 5 μm, (G and H) 0.5 μm, (I) 0.2 μm

### 2. Ultrastructure of PD at the graft interface

We took advantage of our method of CLEM and the accurate targeting of the graft interface to study the PD formed between the scion and rootstock. Plasmodesmata were already present from 3 DAG and every PD observed contained a desmotubule structure in their centre and often had a constricted neck region where the plasma membrane and desmotubule were in very close interaction (Fig. 3). We defined four classes of PDs based on their shapes (Fig. 3A). Class I PD were found in 32.4 % of cases and correspond to isolated straight, simple stranded PD crossing the cell wall. The class II was characterized by tortuous PD containing from three to six branches with compressed neck regions. These branched PD represented 20 % of observed PD. Class III PD corresponds to twinned PD, which were defined as two straight, simple stranded PD (class I) closer to each other than their average length; this class of PD was rarely observed (7.6 %). Unlike classes I-III, class IV, which corresponded to the most frequently found PD at the graft interface (39.4 %), showed the particularity of not spanning the entire cell wall separating the scion and rootstock (Fig. 3E, F). These PD, called hemi-PD, were often seen forming loop-like structures with at least two cytosolic ends opening within the same cell. On two occasions, we observed an unprecedented desmotubule with a “ball of wool”-like structure where compressed ER had rolled onto itself in a huge central cavity (Fig. 3F). We interpreted hemi-PD as failed attempts at secondary PD formation. A similar frequency of hemi-PD was observed on the scion and rootstock sides of the graft interface indicating that the scion and rootstock are similarly able to insert PD at the graft interface. Thus, the cell machinery for the ER maturation in desmotubules and the cell wall modifications seem to be functional in both grafted partners.

**Figure 3.**
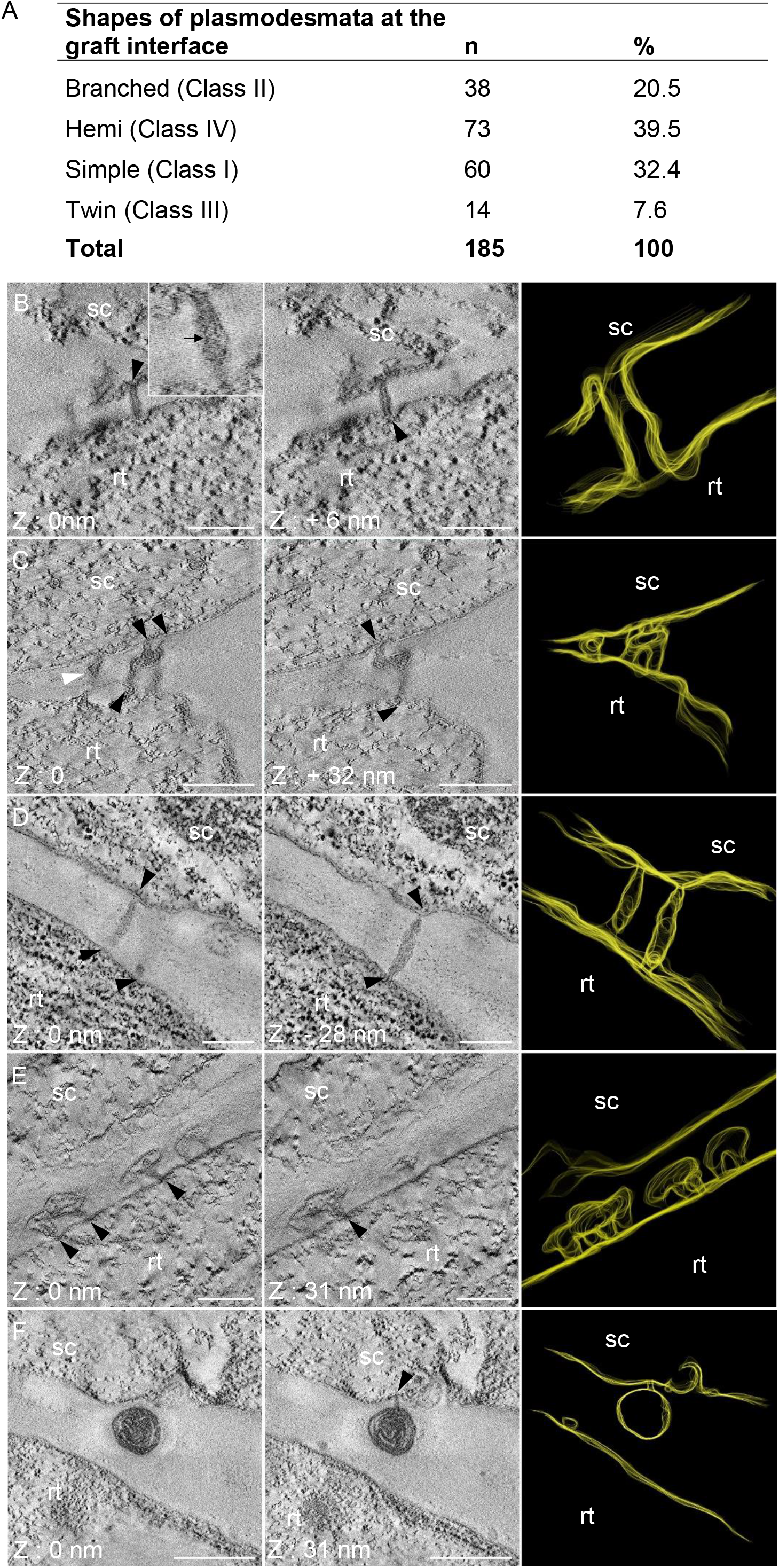
Electron tomograms and three dimensional (3D) segmentation of simple, branched and twin plasmodesmata (PD) **(A)** Number and proportion of the different shapes of PD at the graft interface. **(B-F)** The scion (sc) and the rootstock (rt) are at the top and bottom positions respectively. Black arrowheads pinpoint the extremities of simple **(B)**, branched **(C)** and twin **(D)** PD. For each shape of PD, two Z plans and one 3D segmentation are presented. **(B)** Simple PD with a visible desmotubule (black arrow). **(C)** Branched PD with three branches on the scion side and two branches on the rootstock side, and another PD in its vicinity (B, filled white arrowhead). **(D)** Twin PD characterized by the close proximity of two simple PD. **(E)** Hemi-PD do not span the cell wall and have a large central cavity. **(F)** Hemi-PD of ball of wool presents an aberrant accumulation of desmotubule in this large central cavity. Scale bars: (B, C, D, E) 0.1 μm (F) 0.5 μm

In order to identify events leading to successful symplastic connections between the scion and rootstock, hemi-PD (class IV) and PD of class I, II and III were compared. The measurements of the thickness of the cell wall around 184 PD (60 simple, 38 branched, 14 twin and 72 hemi) revealed that the median of the cell wall thickness around class IV was more 2 times thicker (261 nm) than around PD that crossed the cell wall (115 nm) (Fig. 4). However, no significant difference was found between the classes I to III that successfully made symplastic connections between the scion and rootstock (Fig S3). Interestingly, the median of the penetration depth of hemi-PD into the cell wall was equivalent (99 nm) to the cell wall thickness around class I to III (Fig. 4). Thus, a thin cell wall around 100 nm at the graft interface could be a prerequisite for the establishment of cell-to-cell connections between the scion and the rootstock.

**Figure 4.**
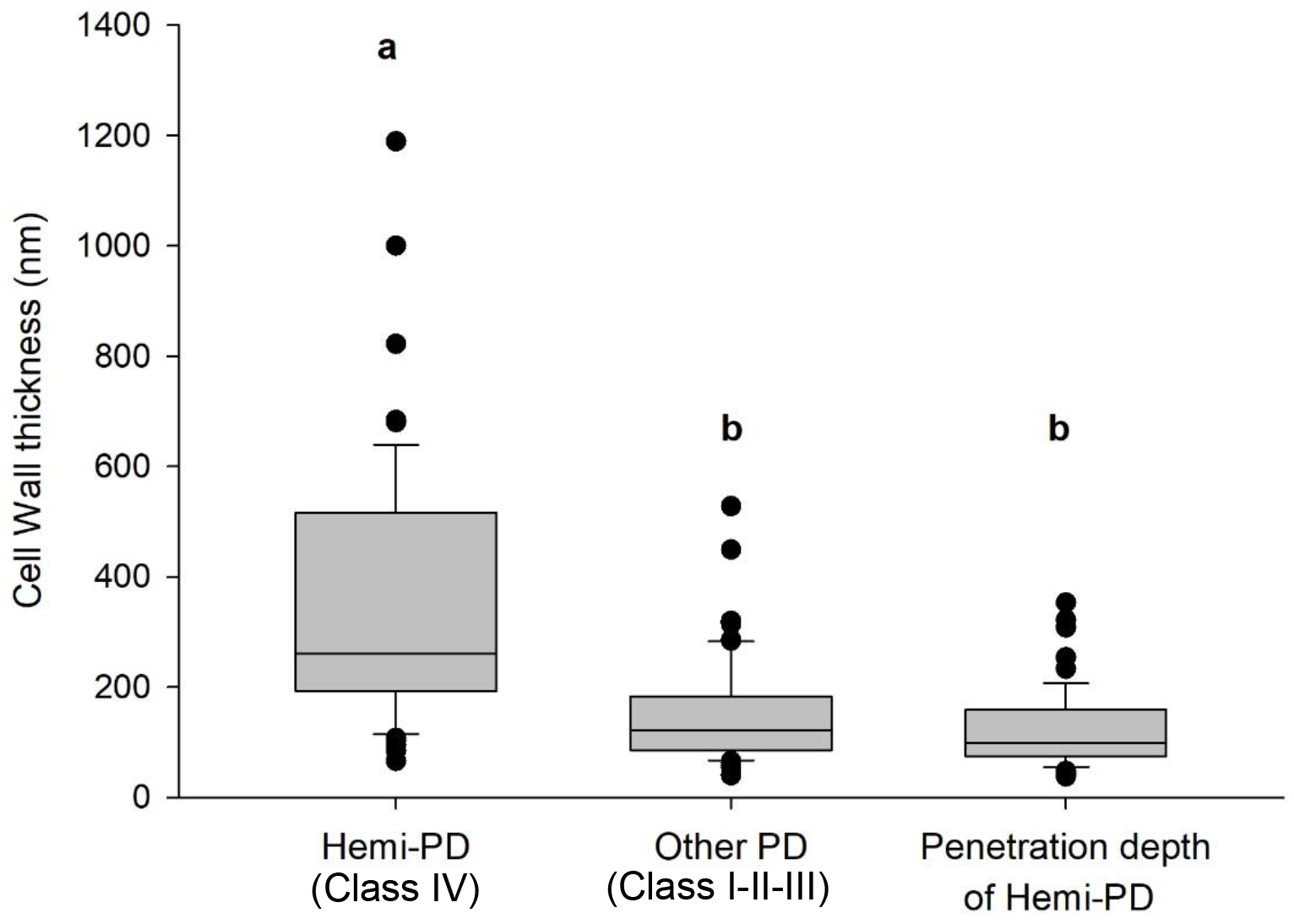
Box plot of the thickness of the cell wall around different classes of plasmodesmata (PD) vs. penetation depth of Hemi-PD at the graft interface. Box plots showing interquartiles and outliers. Letters indicate the results of ANOVA on ranks and Dunn’s tests (p-value < 0.05).

In order to catch PD biogenesis events, we decided to focus on regions where the cell wall is thin. Electron tomography allowed us to reveal ultrastructural details invisible using conventional 2D TEM projections. Electron tomography showed that the cell wall thickness at the graft interface could be reduced down to six nm causing both plasma membranes to be extremely close together (Fig. 5A, B and C). Exceptionally, in one case of extreme cell wall thinning, we catch an event of organelle transfer from one cell to another through a cell wall gap (Fig S4). In this particular graft interface we also found ER-YFP and ER-RFP in the same cells (Fig S5). The regions of extreme cell wall thinning occurred at sites where the plasma membrane and ER were in close contacts (Fig. 5D, E and F), suggesting that it could be a preparation process for a PD insertion by one of the graft partners. Moreover, in regions of thin cell walls of 50 nm, we observed tortuous simple PD spanning the entire cell wall (Fig. 5G, H and I). This suggests that the involvement of only one partner could allow the formation of class I PD. On the contrary, in a thin cell wall of 25 nm, we observed a complex PD in a thicker cell wall spot of 113 nm. This complex PD had 4 arms of scion side and 1 arm on the rootstock side, which met together at a central cavity. This complex PD associated with the local cell wall thickening could be the consequence of the meeting of two PD coming from both partners (Fig. 5J, K and L). In a neighbouring cell, in a thicker cell wall of 134 nm, a hemi-PD was positioned in front of two hemi-PD coming from the other partner (Fig. 5M, N and O). Thus, in this case, both grafted partners inserted PD arms in the same site of the cell wall. Unfortunately, it is impossible to follow the ultrastructure developments of the same PD over time could so we cannot know whether these different arms would fuse together or not. We hypothesize that the large volume of the central cavity of hemi-PD could facilitate hemi-PD contact and fusion across the graft interface.

**Figure 5.**
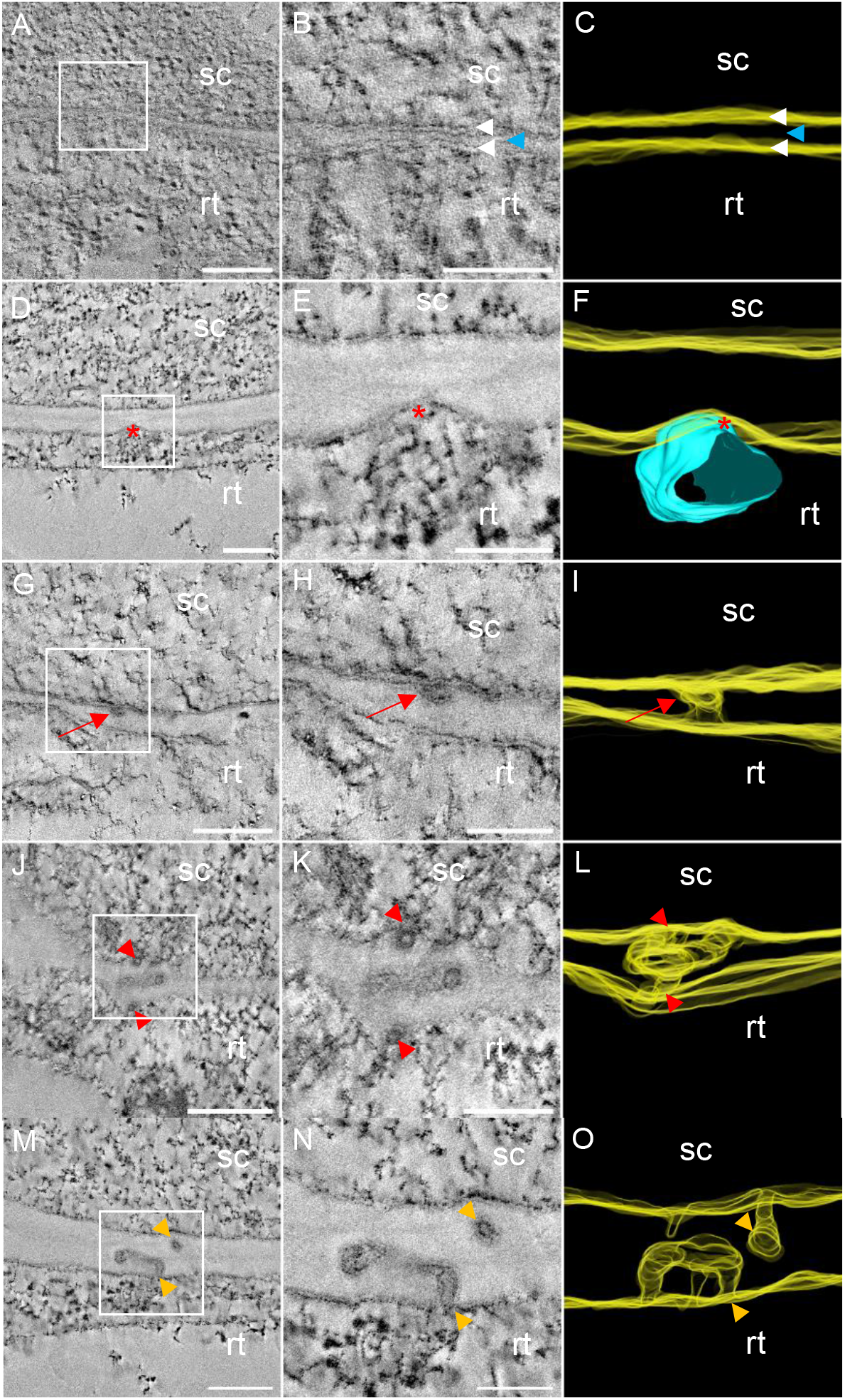
Electron tomograms and 3-D segmentation of plasmodesmata (PD) biogenesis events at the graft interface. The scion (sc) and the rootstock (rt) are respectively at the top and bottom positions. **(A, B, C)** Extreme thinning observed all along the cell wall between two grafted cells. White arrowheads indicate the bilayer of plasma membranes and the blue arrowhead indicates the residual cell wall between them **(D, E, F)** Localized thinning of the cell wall at the endoplasmic reticulum/plasma membrane contact site at the rootstock side (red asterisks). **(G, H, I)** Simple PD (red arrows) observed through a thin cell wall. **(J, K, L)** Complex PD in a localized thickening of a thin cell wall (entrances are depicted with red arrowheads). **(M, N, O)** Hemi-PD facing PD arms at the graft interface (Orange arrowheads). Scale bars: (A, D, G, J, M) 0.2 μm, (B, E, H, K, N) 0.1 μm

Secondary PD formation at the graft interface seems require a cell wall remodelling with a thinning of the cell wall. In addition, the PD characteristics described here could indicate that PD could be formed by a mono-directional PD penetration in the cell wall in cases where the cell wall is thinner than 100 nm. In the cases of thicker cell walls, PD formation could depend on a synchronized spatial-temporal hemi-PD penetration from both graft partners.

## Discussion

Understanding the cellular events occurring during graft union formation has long been a goal for plant scientists. The emergent high-end microscopy technique, CLEM, allowed us to target unambiguously the graft interface and characterize its ultrastructure for the first time. This method allowed us to characterize different events occurring at the graft interface from 3 DAG such as cell differentiation, cell wall modifications, the establishment of communication between the scion and rootstock, and especially to determine the 3D structure of secondary PD formed at the graft interface.

Thanks to the absence of cell-to-cell diffusion of HDEL motif, fluorescent labelling of ER lumen of scions and rootstocks allowed the accurate graft interface. During sample preparation for the TEM, despite the higher brightness of YFP (44.89 unites) than the mRFP one (12.25 unites), the mRFP signal was better preserved during sample processing than YFP. Our hypothesis to explain this difference is that in the acid resin environment (pH_HM20_ = 5), the fluorescent state of mRFP (i.e. the deprotonated state), is better maintained than the fluorescent state of YFP due to differences of their pKa (PkamRFP = 4,5; PkaYFP = 6,9); as has been shown for an autophagic flux probe (Kaizuka et al., 2016). Therefore, choosing FPs with low pKa could be an important parameter to take into account before starting a CLEM approach. Our method allows the preservation of both fluorescence and ultrastructure with a high resolution of around 3 nm, as such this CLEM method could be used to study the ultrastructural events relevant to other biological questions.

In *A. thaliana* homografts, we show that secondary PD are formed at the graft interface by at least 3 DAG (the first time point we were able to study for technical reasons). We distinguished four classes of PD: simple (I), branched (II), twin (III) and hemi (IV). The latter, hemi-PD, have the particularity of not entirely spanning the cell wall and could correspond to failed attempts of PD formation. Hemi-PD were the class the most frequently found at the graft interface, therefore it seems that even between cells of the same species, secondary PD biogenesis at the graft interface appears not to be systematically a successful event. Hemi-PD are found in same proportions on the scion and rootstock sides of the graft interface demonstrating that the scion and the rootstock have the same capacity to insert PD into the cell wall.

Every class of PD had a desmotubule in the central part, which could be simple, branched or forming a loop to a “ball of wool” shape. This study is the first to observe ball of wool shaped PD. The formation of ball of wool shaped PD could be the consequence of hemi-PD having only one cytosolic opening, through which the desmotubule is sent, without the possibility of exit. This suggests that, PD progression into the cell wall is a directed, active mechanism where the desmotubule-plasma membrane interactions observed at the entrances of PD could play a central role. In addition, the accumulation of compressed desmotubule in the central cavity shows that the desmotubule compression is, at least sometimes, independent of direct plasma membrane-ER interactions.

The idea that cell wall modifications at the graft interface are required for the correct establishment of the graft union has been suggested previously (Jeffree and Yeoman, 1983; Kollmann and Glockmann, 1991; Notaguchi et al., 2020). At the graft interface, extreme cell wall thinning was observed; in rare cases this thinning was so excessive that it lead to the formation of gaps in the cell wall and this could explained the presence of some cells that contain both red and green ER and nuclei in our experiments. The idea that gaps form between the cells of the scion and the rootstock is supported by the observation of local horizontal transfer of organelles between the scion and rootstock of grafted plants (Stegemann and Bock, 2009; Stegemann et al., 2012; Lu et al., 2017; Hertle et al., 2021). Cell wall remodeling could be essential to the establishing symplastic connections between the rootstock and scion, and cell wall remodeling could be particularly concentrated at the ER-plasma membrane contact sites. In fact, our results suggest that the PD progression through the cell wall is impaired when the cell wall is much thicker than 100 nm. This could be because of the loss of direct contact between the desmotubule and the plasma membrane at the middle part of cell walls, which have not undergone adequate thinning. In the absence of contacts between the desmotubule and the plasma membrane, the desmotubule could lose its sense of direction and could form a loop to come back to its “home-cell” (hemi-PD), or get lost in a central cavity (ball of wool shaped PD). Thus, the contact site between the ER and the plasma membrane could be a key event in the process of the secondary PD biogenesis by participating in the targeted thinning of the cell wall at the PD entry site and by driving the desmotuble across the cell wall to the neighboring cell. The formation of secondary PD that span the cell wall entirely seems to require cell wall thinning and ER-plasma membrane tethering sites.

The presence of four different PD classes could be the result in several biogenesis mechanisms or different maturation states of PD. Our results show that in the thin cell wall at the graft interface, new secondary PD can belong to the class I, simple PD, potentially formed by a directed active progression of the desmotubule across the cell wall, these PD resemble *de novo* primary PD (Nicolas et al., 2017). The branched PD (class II) could result from the maturation of simple PD by the trapping of ER strands in continuity with the desmotubule during the cell wall thickening, as seen in the maturation of primary PD (Faulkner et al., 2008). This idea is supported by the fact that newly divided cells have a majority of simple PD and that mature cells have a majority of branched PD (Oparka et al., 1999; Nicolas et al., 2017). However, the presence of branched PD at the graft interface is not clearly associated with a general thickening of the cell wall. Furthermore, ultrastructural examinations indicate that at the graft interface branched PD could be in some cases the result of the fusion of two well-aligned hemi-PD, rather than a maturation process. For branched PD biogenesis at the graft interface, the scion and rootstock cells would have to initiate hemi-PD opposite each other at the same moment and at the same site (with a precision around several hundred nanometers in 3D). In this case, the plasma membranes and the desmotubules of both grafting partners have to fuse in order to establish the symplastic continuity between the scion and the rootstock. This symmetric process, on either side of the cell wall at the graft interface, would require the precise spatial-temporal coordination of processes between the scion and the rootstock. This coordination could be mediated by signals produced during the cell wall remodeling (For reviews: Anderson and Kieber, 2020; Duman et al., 2020) and/or by local ER-plasma membrane interactions (For review: Li et al., 2021a). The genesis of a large central cavity, by for example the formation of hemi-PD running in parallel to the cell wall, could increase the chance of alignment of both hemi-PD on either side of the graft interface. If the cell wall is too thick, or the scion and rootstock are not well coordinated, desmotubule entry into the cell wall by only one of the grafted partners occurs, PD progression into the cell wall is deviated and PD evolve into a loop-form, and finally, a hemi-PD is produced. It is possible that branched PD turn into twin PD (class III) during the cell wall expansion as this mechanisms could permit a relatively efficient increase of the number of PD (Faulkner et al., 2008).

This work provides a new view of cell-to-cell connections through PD: it suggests that desmotubules are more than structural components of PD, but could also be dynamic ER compartments actively moving through the cell wall to make PD connections from cell-to-cell. In the future, our CLEM approach could be further exploited to phenotype mutant lines of *A. thaliana* to identify genes involved in the formation of the graft interface as well as providing an improved understanding of secondary PD biogenesis. Thanks to the excellent preservation of both fluorescence and ultrastructural details of this CLEM method, it could also be used on other plant tissues and biological questions in the future.

## Acknowledgements

The master 2 and the Ph.D. thesis of Clément Chambaud was financed by: the grant “PEPS2015 Super-CLEM-3D”, the grant « Etude des mécanismes de l’union greffon/porte-greffe », 50 % from the INRAE department Plant biology and breeding and 50 % from the Région Nouvelle Aquitaine, France. Further financial support was provided by the Starter grant « Etude des mécanismes de l’union greffon/porte-greffe » from the INRAE department Plant biology and breeding. All sample preparation and imaging were done on the Pôle Imagerie du Végétale, appended to the Bordeaux Imaging Centre (http://www.bic.u-bordeaux.fr/) which is a part of the France BioImaging Infrastructure (https://france-bioimaging.org/). The Region Aquitaine also supported the acquisition of the electron microscope (grant No. 2011 13 04 007 PFM). The “Contrat-Plan-Etat-Region PMUSCIVI-Plateforme mutualisée en sciences du vivant 2015_2020” provides the founding for the acquisition of the confocal microscope. Claire Brehelin gave transgenic lines. Thanks respectively to Christophe Garcion and Charles Melnyk to give us access to the plant culture room and show us the grafting procedure.

